# Protein synthesis inhibition and loss of homeostatic functions in astrocytes from an Alzheimer’s disease mouse model: a role for ER-mitochondria interaction

**DOI:** 10.1101/2022.03.24.485644

**Authors:** Laura Tapella, Giulia Dematteis, Marianna Moro, Beatrice Pistolato, Elisa Tonelli, Virginia Vita Vanella, Daniele Giustina, Aleida La Forgia, Elena Restelli, Elettra Barberis, Tito Cali, Marisa Brini, Salvatore Villani, Erika Del Grosso, Mariagrazia Grilli, Marcello Manfredi, Marco Corazzari, Ambra A Grolla, Armando A Genazzani, Dmitry Lim

**Author notes:** These Authors contributed equally (Laura Tapella, Giulia Dematteis). Correspondence should be sent to; Armando A Genazzani and Dmitry Lim.

## Abstract

Deregulation of protein synthesis and ER stress/unfolded protein response (ER stress/UPR) have been reported in astrocytes. However, the relationships between protein synthesis deregulation and ER stress/UPR, as well as their role in the altered homeostatic support of Alzheimer’s disease (AD) astrocytes remain poorly understood. Previously, we reported that in astrocytic cell lines from 3xTg-AD mice (3Tg-iAstro) protein synthesis was impaired and ER-mitochondria distance was reduced. Here we show that impaired protein synthesis in 3Tg-iAstro is associated with an increase of p-eIF2α and downregulation of GADD34. Although mRNA levels of ER stress/UPR markers were increased two-three-fold, we found neither activation of PERK nor downstream induction of ATF4 protein. Strikingly, the overexpression of a synthetic ER-mitochondrial linker (EML) resulted in a reduced protein synthesis and augmented p-eIF2α without any effect on ER stress/UPR marker genes. *In vivo*, in hippocampi of 3xTg-AD mice, reduced protein synthesis, increased p-eIF2α and downregulated GADD34 protein were found, while no increase of p-PERK or ATF4 proteins was observed, suggesting that in AD astrocytes, both *in vitro* and *in vivo*, phosphorylation of eIF2α and impairment of protein synthesis are PERK-independent. Next, we investigated the ability of 3xTg-AD astrocytes to support metabolism and function of other cells of the central nervous system. Astrocyte conditioned medium (ACM) from 3Tg-iAstro cells significantly reduced protein synthesis rate in primary hippocampal neurons. When added as a part of pericyte/endothelial cell (EC)/astrocyte 3D co-culture, 3Tg-iAstro, but not WT-iAstro, severely impaired formation and ramification of tubules, the effect, replicated by EML overexpression in WT-iAstro cells. Finally, a chemical chaperone 4-phenylbutyric acid (4-PBA) rescued protein synthesis, p-eIF2α levels in 3Tg-iAstro cells and tubulogenesis in pericyte/EC/3Tg-iAstro co-culture. Collectively, our results suggest that a PERK-independent, p-eIF2α-associated impairment of protein synthesis compromises astrocytic homeostatic functions, and this may be caused by the altered ER-mitochondria interaction.

## INTRODUCTION

Early cellular dysfunction during AD pathogenesis includes deregulation of Ca^2+^ homeostasis, mitochondrial dysfunction and bioenergetic deficit, oxidative stress and altered cell-cell communication. Such alterations may be traced back to the deregulation of protein synthesis, associated with the activation of endoplasmic reticulum (ER) stress/unfolded protein response (UPR), proposed as targets for the development of AD therapy ^1–3^. Activation of ER stress/UPR has been reported in patients with advanced AD stages ^4–7^. In cellular and animal AD models, heterogeneous and somewhat contrasting data have been reported and activation of ER stress/UPR in AD models has been debated ^8^. The central element which links ER stress/UPR to the accumulation of misfolded proteins is represented by PRKR-like endoplasmic reticulum kinase (PERK)-dependent phosphorylation of eukaryotic initiation factor 2α (eIF2α). In turn, this protein inhibits assembly of ribosomal 43S preinitiation complex and imposes a global translational block, with a profound impact on neural cell physiology ^9,10^. However, non-canonical variants of ER stress/UPR and their role in AD pathogenesis have been discussed ^11,12^. While most of the reports consider neuronal mechanisms of ER stress/UPR in AD, contribution of astrocytes has been generally overlooked.

Astrocytes are homeostatic and secretory cells, whose function is to warrant the stability of the extracellular space, the development and correct functional integration of the CNS components in an environment which has recently been called the “active milieu” ^13^. Therefore, the activation of ER stress/UPR and deregulation of protein synthesis in astrocytes would be particularly important for their potential role in CNS pathologies in terms of cellular dysfunction and loss of supportive functions. For example, local translation of mRNA in astrocytic processes is suggested to be important for shaping the repertoire of astrocytic plasma membrane and secreted proteins warranting support to neurons ^14,15^. A derangement of ribosomal protein synthesis machinery in AD astrocytes has already been documented ^4^. While a canonical [PERK → p-eIF2α → protein synthesis block] pathway is postulated in astrocytes, only fragmentary data are available. Moreover the relationships between ER stress/UPR and protein synthesis in astrocytes during AD progression remain largely unexplored^16^.

Recently, we proposed immortalized hippocampal astrocytes from 3xTg-AD mice (3Tg-iAstro cells) as a novel cellular model which shows features of AD-like pathology, i.e., transcriptional and translations alterations, deregulation of Ca^2+^ signaling, bioenergetic deficit, elevated ROS and augmented ER-mitochondria interaction ^17–21^. The central finding, linking the astrocytic cell pathology with possible deficit of homeostatic support, was protein synthesis impairment and a modest increase of ER stress/UPR related genes ^19,20^. Herein we further investigated, both *in vitro* and *in vivo*, if reduction of protein synthesis in 3xTg-AD astrocytes was due to ER stress/UPR. Our results suggest that a PERK-independent, p-eIF2α-associated impairment of protein synthesis alters secretome of AD astrocytes and compromises their supportive and defensive functions, possibly through altered ER-mitochondria interaction.

## RESULTS

### Protein synthesis impairment in AD astrocytes is associated with PERK-independent phosphorylation of eIF2α in 3Tg-iAstro cells

To investigate if protein synthesis impairment in 3Tg-iAstro cells was due to activation of [PERK → eIF2α → activating transcription factor 4 (ATF4)] axis, we first of all confirmed that 3Tg-iAstro have a significant reduction of protein synthesis, compared to WT-iAstro cells, using both immunocytochemical (ICC) (Fig. 1a) and Western blot (WB) analysis (Fig. 1b) of puromycin incorporation in neo-synthetized peptides (SUrface SEnsing of Translation (SUnSET) method ^22,23^) (Fig. 1). Next, we investigated expression levels of p-eIF2α, whose de-phosphorylation is essential for the assembly of pre-initiation complex and initiation of translation ^10^. As shown in Fig. 2a, the levels of p-eIF2α were significantly higher in 3Tg-iAstro compared with WT-iAstro cells, and comparable to levels in WT-iAstro and 3Tg-iAstro treated with thapsigargin (THG), an established ER stress/UPR inducer. We also measured expression levels of growth arrest and DNA damage-inducible gene 34 GADD34, a protein which provides a scaffold for eIF2α de-phosphorylation by protein phosphatase 1 (PP1) ^9^. Surprisingly, GADD34 protein levels were significantly lower in 3Tg-iAstro compared to WT-iAstro cells (Fig. 2a). Next, we asked if augmented levels of p-eIF2α correlated with activation of PERK in our cellular model. However, the levels of p-PERK were not different in WT-iAstro and 3Tg-iAstro cells (Fig. 2b). During ER stress/UPR, activated and auto-phosphorylated PERK phosphorylates eIF2α and induces p-eIF2α-dependent upregulation of transcription factor ATF4 ^9^. However, in 3Tg-iAstro levels of ATF4 were not different from those in WT-iAstro cells (Fig. 2b). Altogether these data suggested that phosphorylation of eIF2α in 3Tg-iAstro cells was not due to activation of [PERK → eIF2α/GADD34 → ATF4] axis. These results are in apparent contrast with our previous report of a two-three-fold transcriptional upregulation of ER stress/UPR-induced genes Atf4, spliced variant of X-box-binding protein 1 (Xbp1s) and homocysteine inducible ER protein with ubiquitin like domain 1 (Herpud1) in 3Tg-iAstro cells compared to its WT counterpart ^20^. Therefore, we compared the induction of ER stress/UPR markers in 3Tg-iAstro cells with those induced by THG, which produces maximal induction of ER stress/UPR markers. We confirmed that in 3Tg-iAstro cells mRNA of the three ER stress/UPR markers significantly increased, and the increase was in line with our previous publications ^20^ (Fig. 3, middle histograms). However, the maximal upregulation of all three transcripts was much higher in THG-treated (1 μM, 4 h) vs non-treated cells than that in 3Tg-iAstro vs WT-iAstro (by 4.8 fold for Atf4, by 50-70 fold for Xbp1s and 18-25 fold for Herpud1) (Fig. 3, right histograms). Of note, there was a tendency to a lower THG-induced upregulation of Xbp1s and Herpud1 (mRNA, Fig. 3b,c right histograms), and ATF4 (protein, Fig. 2b) in 3Tg-iAstro compared with WT-iAstro, although the differences were not significative in the current experimental setting. To strengthen the result we assessed mRNA levels of other genes involved in different arms of ER stress/UPR response as well as UPR-inducible chaperons including Atf6, Ddit3/CHOP, Bip/Grp78, calreticulin and Dnajb9/ERdj4. As shown in Supplemental Fig. 1, essentially the same result was obtained. We concluded that eIF2α phosphorylation and the reduction of protein synthesis in 3Tg-iAstro cells were PERK-independent, however a low-grade chronic ER stress cannot be ruled out.

**Figure 1.**
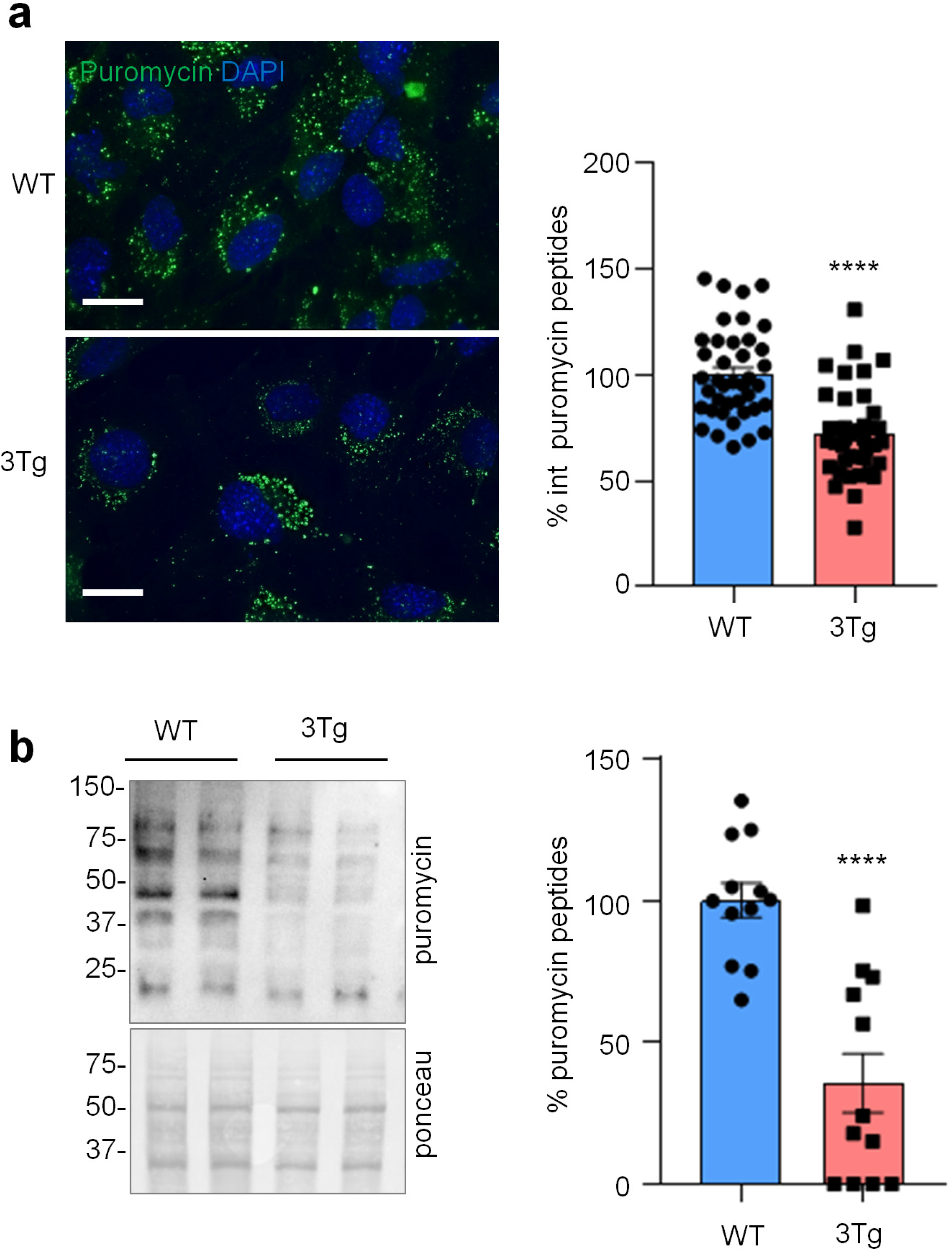
Protein synthesis analysis in WT- and 3Tg-iAstro cells. **(a)** Cells were pulsed with puromycin (4 μM, 1.5 h), fixed and analysed by IF with anti-puromycin antibody (green) and with DAPI to stain nuclei (blue). Images were acquired with Leica Thunder imager 3D live cell microscope, scale bar = 25 μm. Data are expressed as mean ± SEM, WT-iAstro cells n = 40, 3Tg-iAstro cells n = 40, from 4 independent experiments; ****, p < 0.05 by unpaired t test. **(b)** WB with anti-puromycin antibody and ponceau staining on cells treated with 4 μM puromycin. Data are expressed as mean ± SEM of 12 independent experiments; ****, p < 0.0001 by unpaired t test.

**Figure 2.**
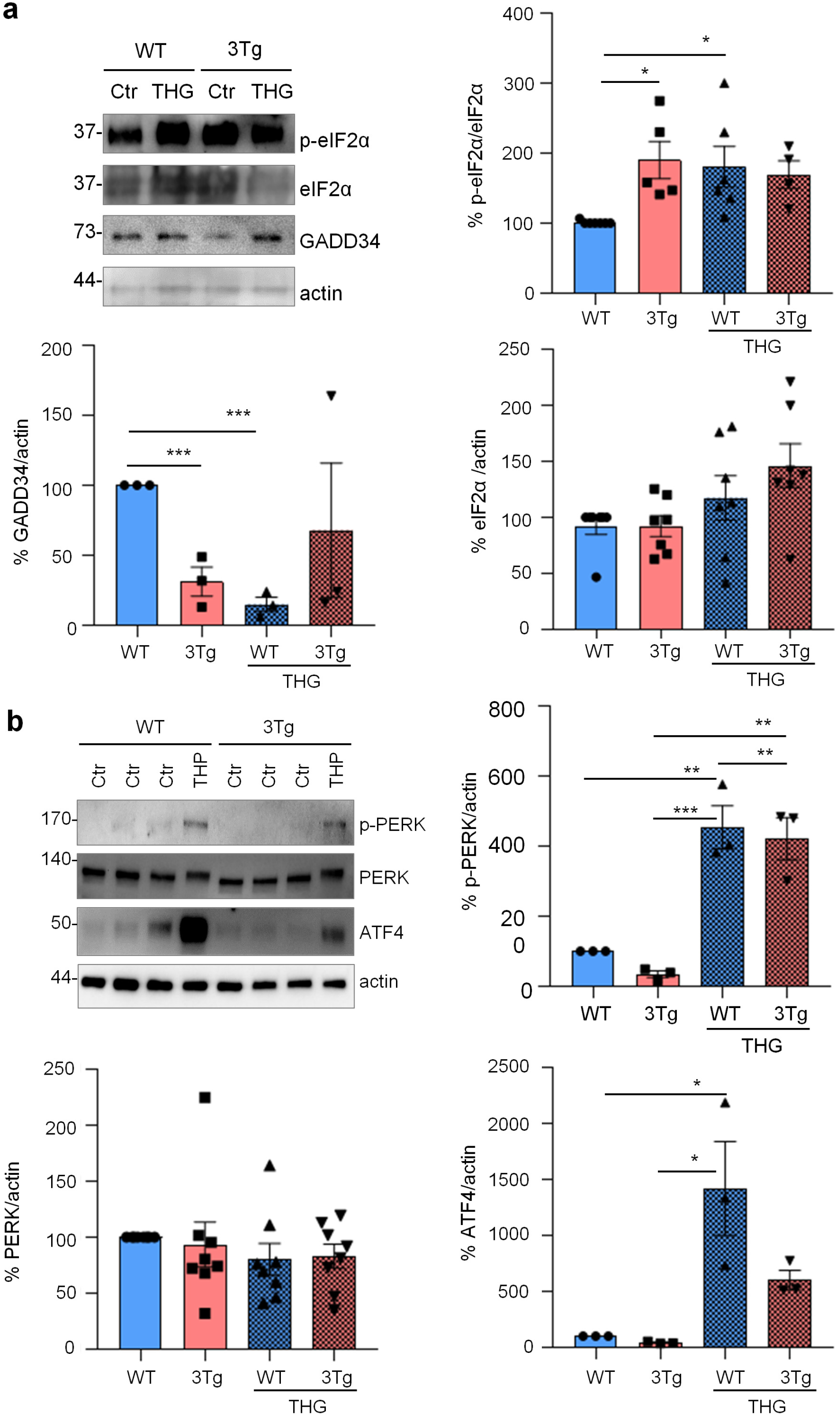
PERK-independent activation of p-eIF2α. **(a)** Analysis of eIF2α phosphorylation and GADD34 expression on WT and 3Tg-iAstro. Cells were treated or not with 1 μM THG for 1 h, lysed and analysed by WB with anti-p-eIF2α, eIF2α, GADD34 and actin antibodies. Data are expressed as mean ± SEM of n = 4 (3Tg-iAstro + THG), n = 5 (3Tg-iAstro) or n = 6 (WT- and WT-iAstro + THG) independent experiments; *, p < 0.05 by one-way ANOVA, Sidak’s multiple comparison; ***, p<0.001 Dunnet’s multiple comparisons. **(b)** Analysis of PERK phosphorylation and ATF4 induction on WT- and 3Tg-iAstro cells. WB analysis of cells treated as in (a) with anti-p-PERK, PERK, ATF4 and actin antibodies. Data are expressed as mean ± SEM from n = 3 (p-PERK and ATF) or n = 8 (PERK) independent experiments. *, p < 0.05, **, p < 0.01 and ***, p < 0.001 by one-way ANOVA, Sidak’s multiple comparison.

**Figure 3.**
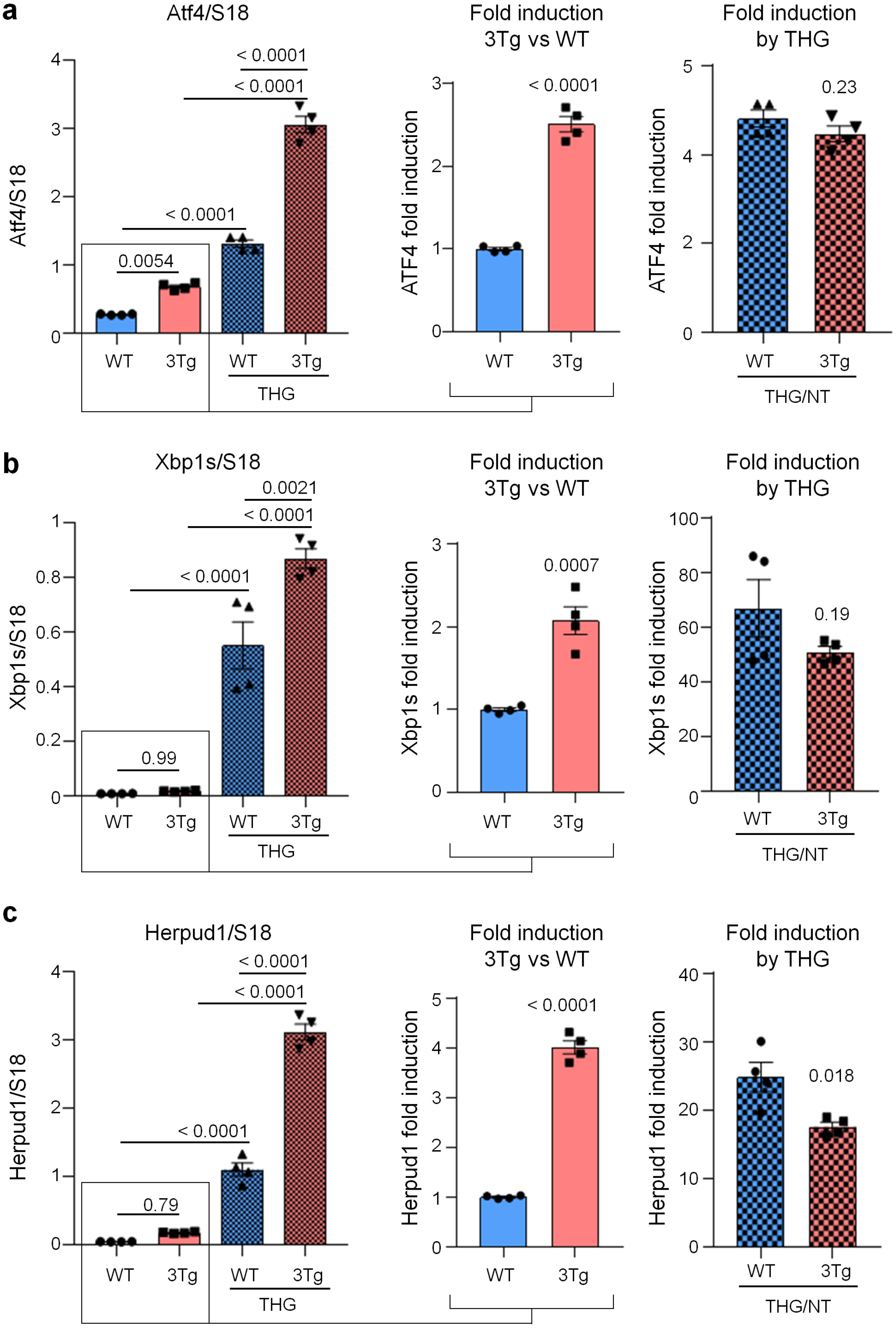
ER stress/UPR genes induction in WT- and 3Tg-iAstro cells. Real-time PCR of Atf4 **(a)**, Xbp1s **(b)** and Herpud1 **(c)** transcripts in cells treated or not with 1 μM THG for 4h. Data of untreated WT- and 3Tg-iAstro cells (middle plots) and THG treated /untreated cell (right plots) are presented separately. Values represent mean ± SEM ΔC(t) of gene/S18 of 4 independent experiments for each condition. Left plots were analyzed using ANOVA, with Tukey posthoc test; middle and right plots were analyzed using unpaired two-tail Student’s t-test.

### Reduction of protein synthesis and increase of p-eIF2α in astrocytes expressing a 10 nm ER-mitochondrial linker

Previously, we reported that ER-mitochondrial interaction, measured by SPLICS (split-GFP-based Contact site Sensor), a recently developed ER-mitochondria contact sites sensor ^24–26^, is increased in 3Tg-iAstro compared with WT-iAstro cells, suggesting a correlation between ER-mitochondrial distance and reduction of protein synthesis ^20^. To test the hypothesis of the causal role of the shorter ER-mitochondrial distance on protein synthesis reduction we transfected WT-iAstro cells with a synthetic linker which fixes the ER-mitochondria distance at about 10-12 nm (named as 10nm-EML) (a kind gift from György Csordás and György Hajnóczky, Thomas Jefferson University). The linker was composed of monomeric red fluorescent protein (mRFP) and an amino acidic liner, flanked at the N-terminal side by an ER-targeting sequence, and at the C-terminal side by an outer mitochondrial membrane-targeting sequence ^27^. We found that the expression of 10nm-EML significantly reduced protein synthesis rate as tested in the SUnSET assay, either by WB (Fig. 4a) or ICC (Fig. 4c). Strikingly, 10nm-EML expression also significantly augmented p-eIF2α levels (Fig. 4b). At this point, we checked if the reduction of protein synthesis and increase of p-eIF2α were paralleled by an induction of ER stress/UPR marker genes. However, expression of Atf4, Xbp1s and Herpud1 transcripts were not different between Ctr (WT-iAstro transfected with mRFP) and 10nm-EML-expressing WT-iAstro cells. Altogether these data, in line with alterations found in 3Tg-iAstro cells, suggest that increased interaction between ER and mitochondria augments phosphorylation of eIF2α and reduces protein synthesis by an UPR-independent mechanism.

**Figure 4.**
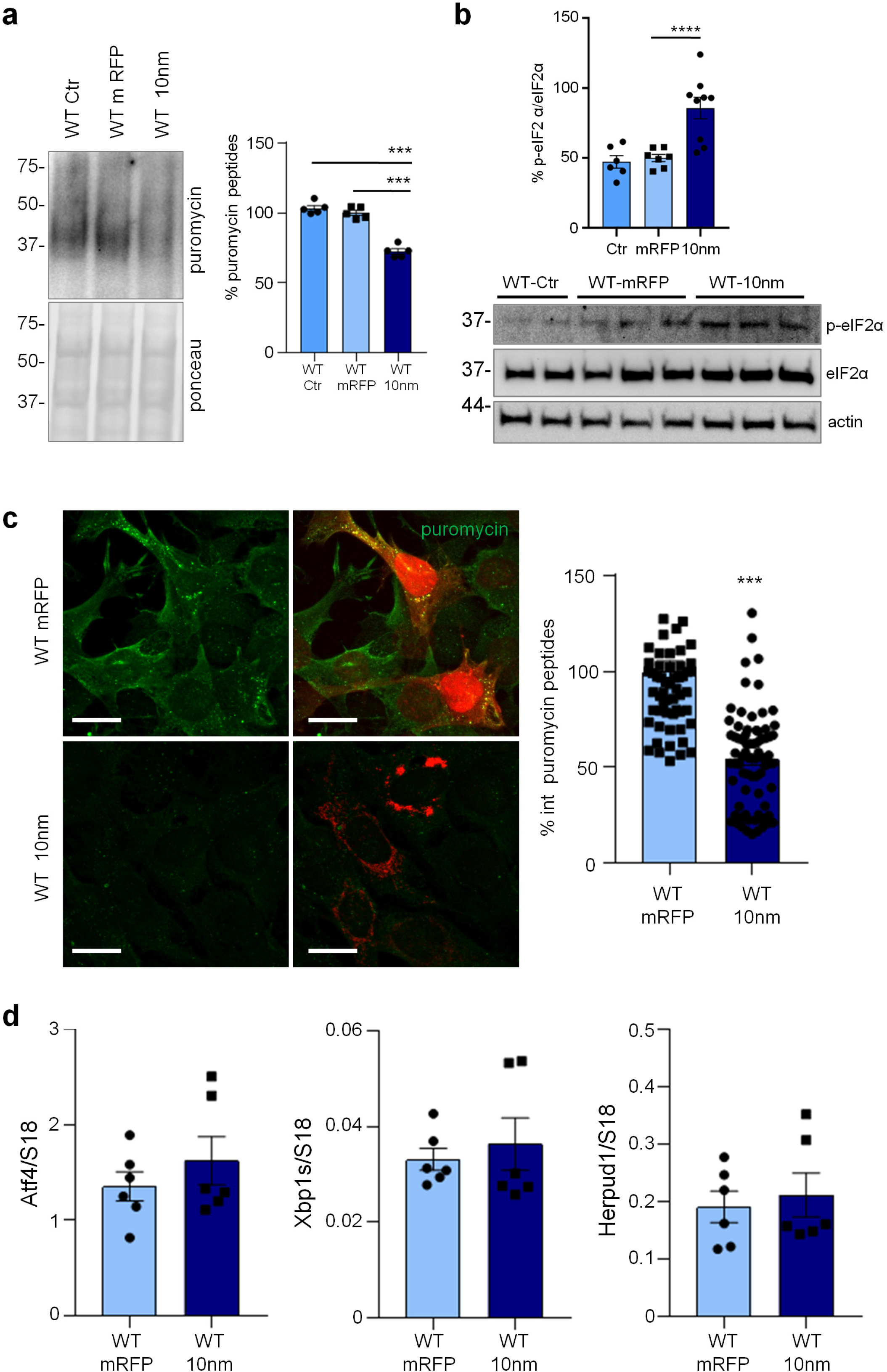
Forced ER-mitochondria interaction causes impairment of protein synthesis in WT-iAstro cells. WT-iAstro cells were non-transfected (WT Ctr) or transfected with mRFP (WT mRFP) and 10-nm ER-mitochondrial linker (WT 10nm). **(a)** WB with anti-puromycin antibody and ponceau staining on lysates of cells treated with 4 μM puromycin; data are expressed as mean ± SEM of 3 independent experiments; ***, p<0.001 by one-way ANOVA, Sidak’s multiple comparison. **(b)** WB analysis of eIF2α phosphorylation. Data are expressed as mean ± SEM of 4 independent experiments; ****, p < 0.0001 by one-way ANOVA, Sidak’s multiple comparison. **(c)** Cells pulsed with 4 μM puromycin were fixed and analysed by IF with anti-puromycin antibody (green). Images were acquired with Zeiss 710 confocal laser scanning microscope, data are expressed as mean ± SEM of 3 independent coverslip; ***, p < 0.001 by unpaired t test. Scale bar = 25 μm. **(d)** Real-time PCR of Atf4, Xbp1s and Herpud1 transcripts in cells transfected with mRFP and 10-nm ER-mitochondrial linker; data are expressed as mean ± SEM of 4 independent wells.

### PERK-independent increase of p-eIF2α and protein synthesis reduction in vivo in 3xTg-AD mouse astrocytes

Next, we assessed if similar alterations in protein synthesis, p-eIF2α and ER stress markers could be found also *in vivo* in the hippocampus of 3xTg-AD and WT mice. To assess protein synthesis rate, 3xTg-AD mice were injected with puromycin (225 mg/Kg body weight) intraperitoneally (i.p.) for 1.5 h. Then hippocampi were harvested and puromycin incorporation was analysed by WB and immunohistochemical analysis (IHC). As shown in Fig. 5a, WB analysis showed a significantly reduced puromicyn-positive signal in 3xTg-AD mice compared with WT. The result was confirmed by anti-puromycin staining of brain cryosections (Fig. 5b). Assessment of [PERK → eIF2α/GADD34 → ATF4] axis activation by WB revealed significant increase of p-eIF2α, reduction of GADD34 protein, while total PERK and ATF4 were not changed. Under the same experimental conditions, p-PERK was undetectable by WB analysis in both genotype samples (Fig. 6a). This result, and the absence of total PERK mobility shift, which accompany THG-induced PERK phosphorylation in astrocytes (Fig. 2b), suggest that PERK is not activated in hippocampi of 3xTg-AD mice. IHC analysis confirmed upregulation of p-eIF2α specifically in CA1 neuropil astrocytes of 3xTg-AD mice (Fig. 6b), while GADD34 staining was diffused and was significantly reduced in the CA1 neuropil (Fig. 6c). qPCR analysis on whole hippocampal lysates revealed no changes in Atf4, Xbp1s, and Herpud1 transcript levels (Fig. 6d). Altogether, these data suggest that in AD astrocytes, both *in vitro* and *in vivo*, p-eIF2α-associated reduction of protein synthesis was independent of PERK activation but may be associated with alterations in ER-mitochondria interaction.

**Figure 5.**
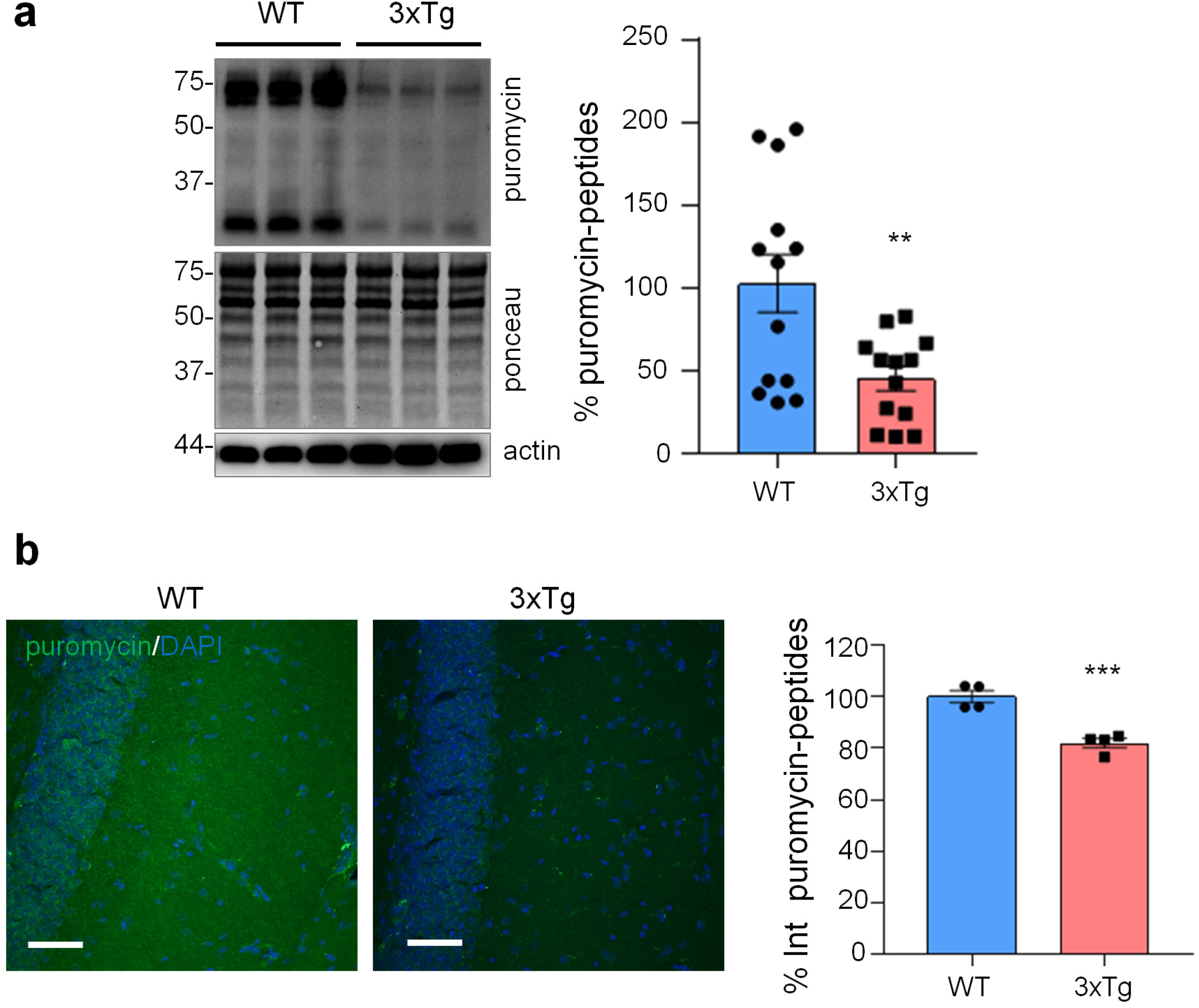
Protein synthesis analysis in WT and 3xTg-AD mice. WT and 3xTg-AD mice were i.p. injected with 225 mg/Kg di puromycin and sacrificed 1.5 h post-injection. **(a)** WB with anti-puromycin, anti-actin antibodies, and ponceau staining on hippocampal homogenates. Data are expressed as mean ± SEM of 13 independent WB from 2 mice per genotype. **, p < 0.01, unpaired t test. **(b)** IF with anti-puromycin antibody (green) and DAPI (blue) of hippocampal brain slices. Data are expressed as mean ± SEM of 4 sections collected from 2 mice per genotype. Scale bar = 50 μm.

**Figure 6.**
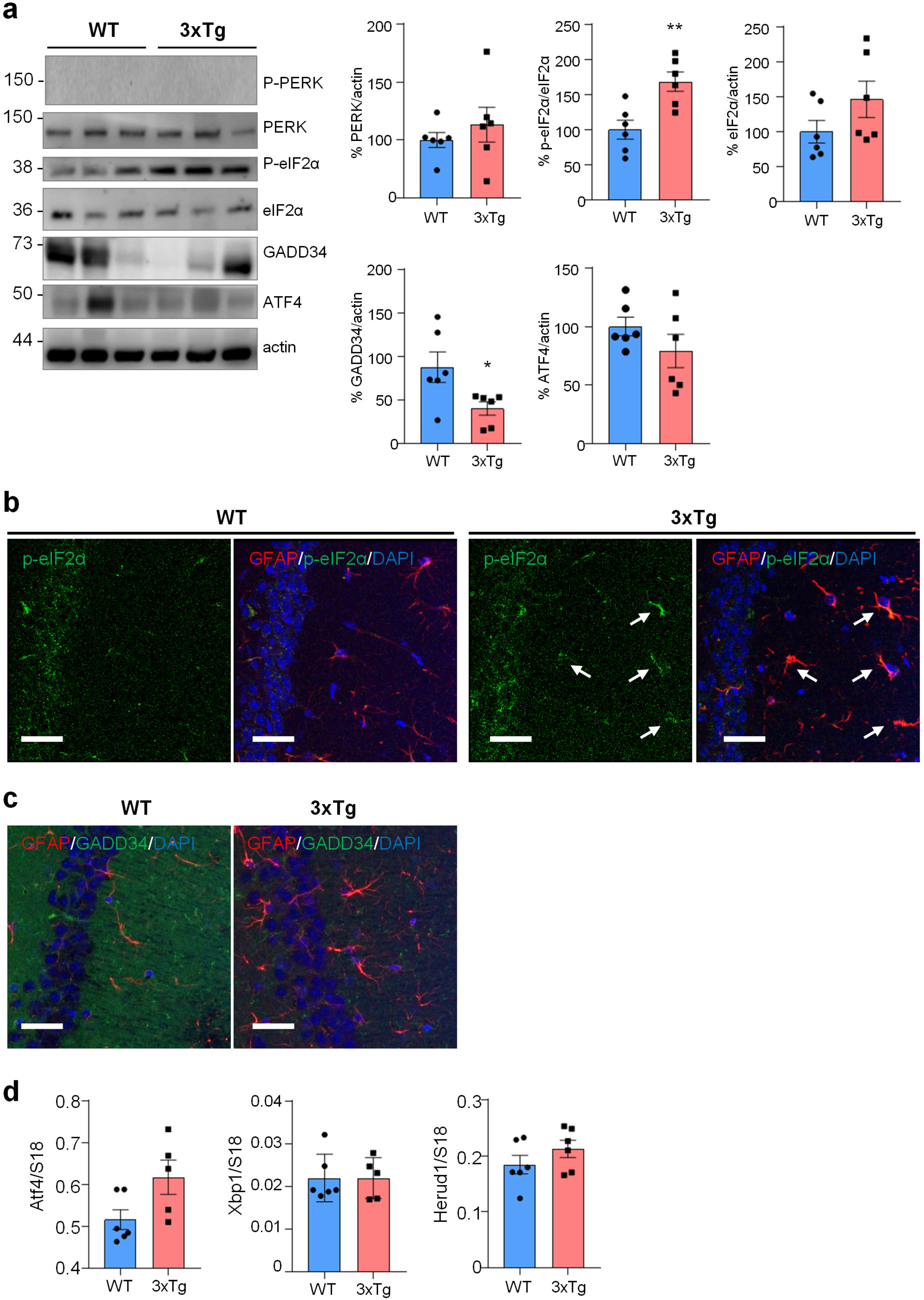
ER stress/UPR pathway in WT and 3xTg-AD mice. **(a)** WB analysis with anti-PERK, p-eIF2α, eIF2α, GADD34, ATF4 and actin on hippocampal homogenates of WT and 3xTg-AD (3xTg) mice. Data are expressed as mean ± SEM of 6 independent experiments; *, p < 0.05; **, p < 0.01 by unpaired t test. **(b, c)** Co-localization of p-eIF2α and GADD34 with GFAP and analysis on WT and 3xTg hippocampi. **(b)** IF on 40 μm thick brain slices with anti-GFAP (red), p-eIF2α (green) and DAPI (blue). Arrows indicate p-eIF2α-expressing GFAP-positive astrocytes. **(c)** IF on 40 μm brain slices with anti-GFAP (red) and GADD34 (green) and DAPI (blue). Images were acquire using Leica SP8 LIGHTNING Confocal Microscope imaging systems, scale bar = 25μm. **(d)** qPCR of Atf4, Xbp1s and Herpud1 on the hippocampi of WT and 3xTg mice. Data are expressed as mean ± SEM of 5-6 independent experiments, unpaired t test analysis.

### 3Tg-iAstro cells do not support neuronal protein synthesis and pericyte-endothelial cell (EC) tubulogenesis in vitro, the effect, replicated by 10nm-EML

Growing body of evidence suggest that non-cell autonomous mechanisms of neuronal degeneration during AD pathogenesis could be mediated by astroglia dysfunction and reduced homeostatic support to neurons and other cells in the CNS. Therefore, we assessed the effect of 3Tg-iAstro cells on neurons and a three-cell pericyte/EC/astrocyte 3D co-culture. 3Tg-iAstro-Conditioned Medium (ACM) transfer, but not WT-iAstro-ACM, onto primary murine hippocampal cultured neurons resulted in a significant reduction of neuronal protein synthesis, an early sign of neuronal dysfunction ^28–31^ (Fig. 7a and b). Treatment of neurons with ACM from WT-iAstro cells overexpressing 10nm-EML had also reduced protein synthesis rate (Fig. 7c). When WT-iAstro or 3Tg-iAstro cells were added as a component of pericyte/EC/astrocyte 3D co-culture, WT-iAstro, but not 3Tg-iAstro, supported formation of vessel-like tubules by pericyte and EC (Fig. 7d). Strikingly, the effect of 3Tg-iAstro was reproduced by co-culture with 10nm-EML-overexpressing WT-iAstro cells (Fig. 7d). These results suggest that alterations of astrocyte-derived soluble factors and cell-cell contact may account for 3Tg-iAstro inability to support neuronal protein synthesis and pericyte/EC tubulogenesis and that ER-mitochondria interaction may have a role.

**Figure 7.**
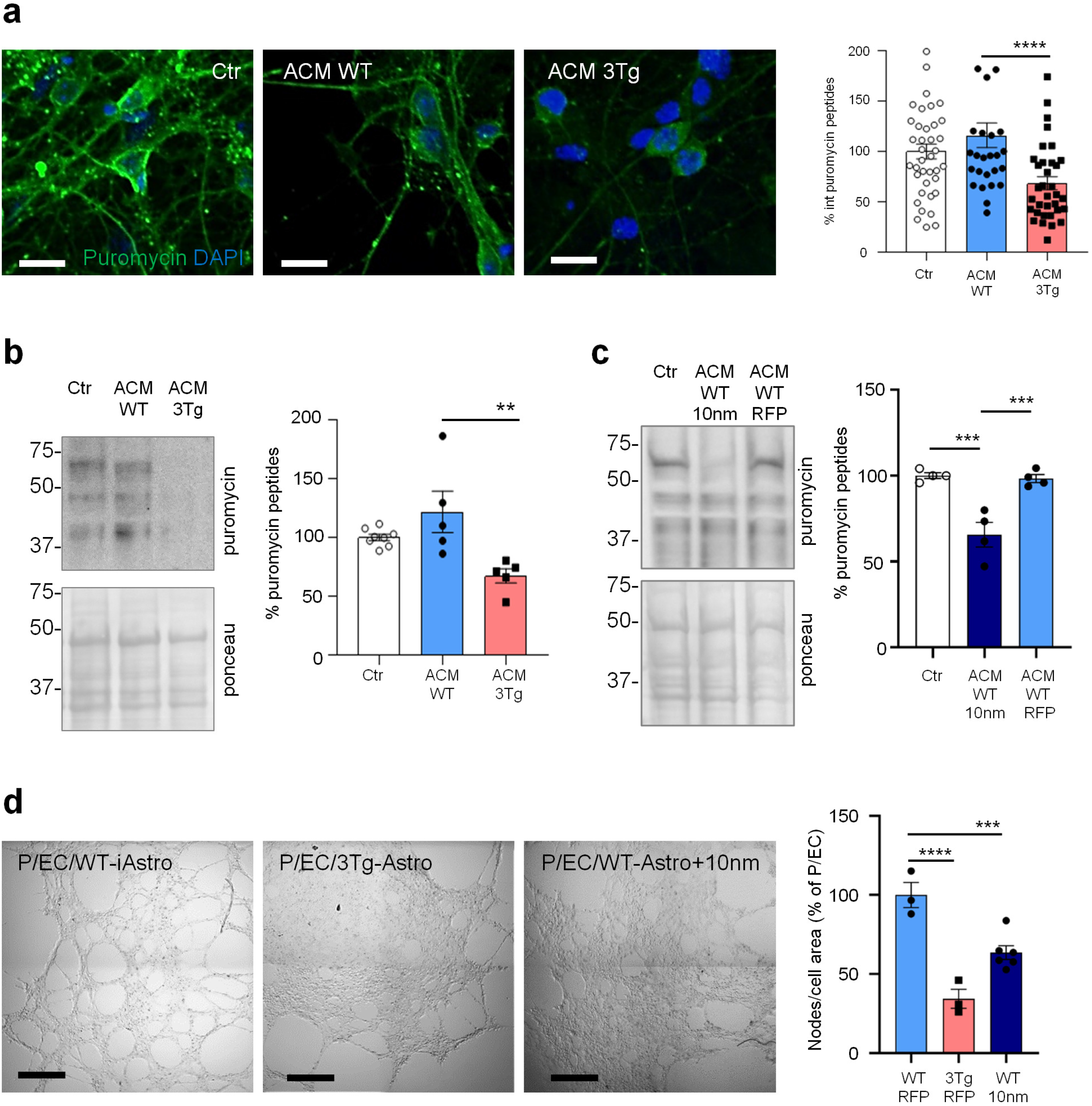
Compromised homeostatic functions of 3Tg-iAstro. **(a)** ACM effects on neuronal protein synthesis. Primary hippocampal neurons were treated with ACM from WT- or 3Tg-iAstro for 6 days (from DIV6 to DIV12) and protein synthesis was evaluated treating cells with 4 μM puromycin for 1 h. IF images of anti-puromycin staining (green) and DAPI (blue) were acquired with FV-1000 Olympus laser confocal scanning system, scale bar = 30 μm. Data are expressed as mean ± SEM from n = 39 Ctr, n = 27 ACM WT-iAstro, n = 34 ACM 3Tg-iAstro from 4 independent cultures; ***, p < 0.001 by one-way ANOVA Sidak’s multiple comparisons. **(b)** WB analysis with anti-puromycin, actin and ponceau staining of neuronal lysates treated as in (a). Data are expressed as mean ± SEM from n = 8 (Ctr) or n = 5 (ACM WT and ACM 3Tg) independent experiments; **, p < 0.01 by one-way ANOVA, Sidak’s multiple comparisons. **(c)** WB analysis with anti-puromycin and ponceau staining of neuronal lysates treated or not with ACM from WT-iAstro or WT-iAstro expressing 10nm-EML. Data are expressed as mean ± SEM of 4 independent experiments; ***, p < 0.001 by one-way ANOVA, Sidak’s multiple comparisons. **(d)** Co-culture of pericytes and endothelial cells (EC) with either WT-iAstro (P/EC/WT-iAstro) or 3Tg-iAstro (P/EC/3Tg-iAstro)) or WT-iAstro expressing the 10nm-EML (P/EC/WT-iAstro+10nm) cells on a layer of Matrigel. Images were acquired by Zeiss 710 confocal laser scanning microscope, scale bar = 500 μm. Data are expressed as mean ± SEM of n = 3 (WT-RFP and 3Tg-RFP) or n = 6 (WT-10nm) independent experiments; ***, p < 0.001, ****, p < 0.0001, by oneway ANOVA Sidak’s multiple comparisons.

### Analysis of secretome from 3Tg-iAstro cells suggests impairment of neurogenic, neuroprotective functions and inter-cellular interaction

In search of astrocyte-derived soluble factors we performed shotgun mass spectrometry proteomics of ACM from WT-iAstro and 3Tg-iAstro cells, followed by bioinformatic analysis. As shown in Supplemental Table 1, 120 and 84 proteins were identified, respectively, in WT- and 3Tg-iAstro cells ACM. Of these, 55 were expressed by both types of astrocytes, while 65 and 29 were identified only in WT- or 3Tg-iAstro cells ACM, respectively. Two pipelines of analysis have been performed. Firstly, proteins were quantified and differentially expressed proteins (DEPs) in 3Tg-iAstro vs WT-iAstro cells were identified (Supplemental Table 2). Five DEPs were identified. Of those, one, fatty acid-binding protein 3 (Fabp3) was upregulated, while four proteins, secreted protein acidic and cysteine rich (SPARC), heat shock protein 90 (HSP90), heat shock protein 73 (HSP73) and α1-tubulin, were significantly down-regulated in 3Tg-iAstro compared with WT-iAstro cells (Fig. 8, left table). SPARC is a pro-neurogenic factor released by astrocytes which promotes neuronal differentiation ^32^. Extracellular heat shock proteins are known to be neuroprotective ^33^. Specifically, extracellular HSP90 protects neurons from oxidative stress ^34^. Therefore, our proteomic results suggest that neurogenic and neuroprotective support in 3Tg-iAstro cells may be reduced compared with the WT counterpart. In a separate analysis, uniquely identified proteins were considered and subjected to gene ontology (GO) analysis using DAVID online tool in search of overrepresented groups of proteins. This analysis revealed that WT-iAstro, but not 3Tg-iAstro secretome, was enriched in proteins involved in cell-cell contacts, focal adhesion contacts and constituents of extracellular matrix (ECM), suggesting that support of cell-cell communication and ECM formation may be impaired in 3Tg-iAstro cells (Fig. 8, right upper table). Next we investigated if manipulation with ER-mitochondrial distance or normalization of protein folding had an effect on the secreted proteins. 10nm-EML overexpression in WT-iAstro cells did not influence the identified proteins (Supplemental Table 3). However, treatment of 3Tg-iAstro cells with a chemical chaperone 4-PBA (4-phenil butyric acid) rescued the presence of proteins responsible for extracellular matrix formation (Fig.8 right bottom table and Supplemental Table 4).

**Figure 8.**
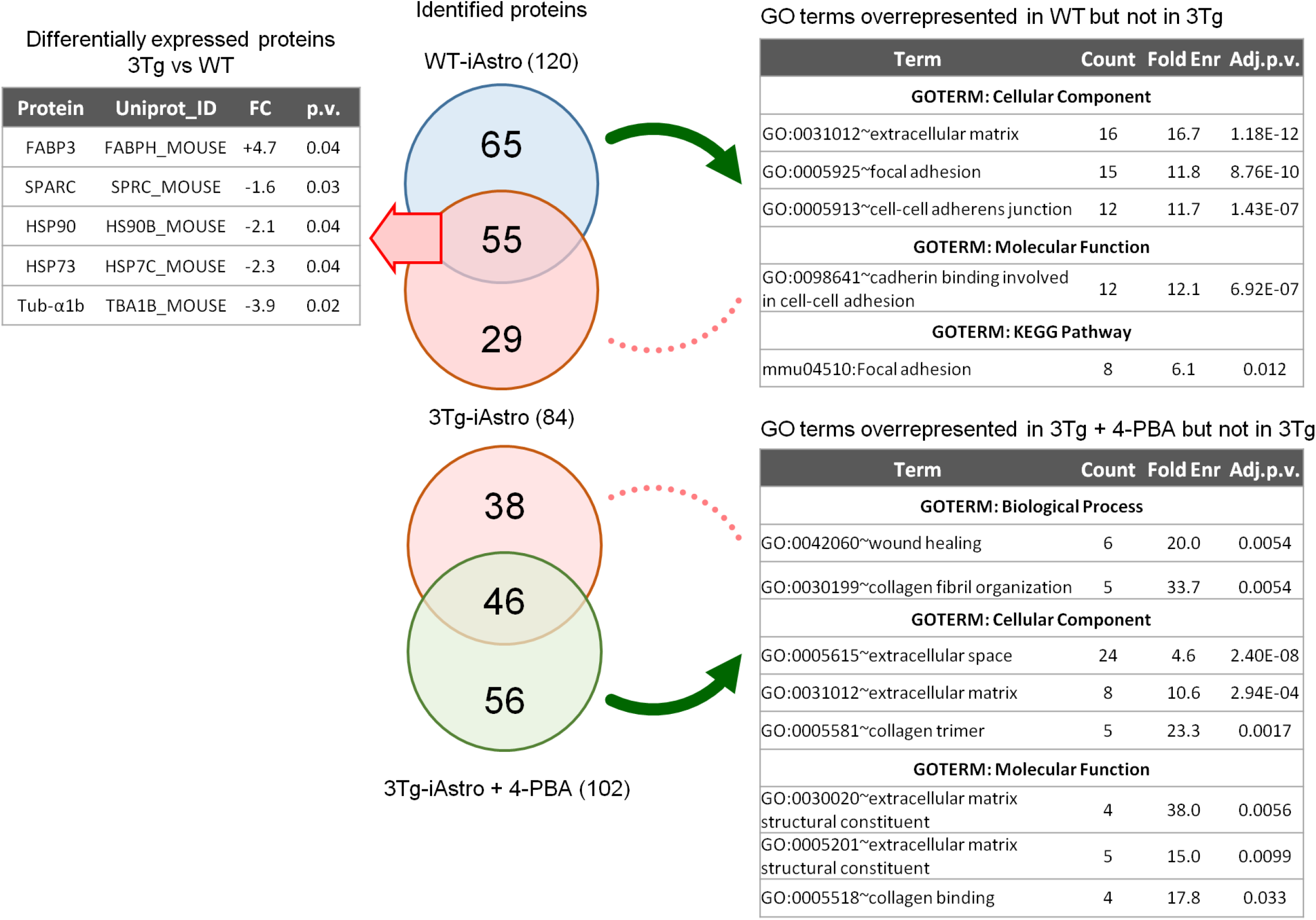
Proteomic analysis of WT- and 3Tg-iAstro secretome. ACM (10 ml) was collected from 48 h culture of WT-iAstro, 3Tg-iAstro and 3Tg-iAstro cells treated with 4-PBA (3 μM, 48h). Proteins were precipitated by TCA and processed as described in Methods section. 120, 84 and 102 proteins were identified in WT-iAstro, 3Tg-iAstro and 3Tg-iAstro + 4-PBA ACM, respectively. Protein quantification of 3Tg-iAstro vs WT-iAstro ACM returned five differentially regulated proteins (1.3 fold change cut-off, p < 0.05) (left upper table). Gene ontology analysis of proteins, unique for WT-iAstro or 3Tg-iAstro + 4-PBA both compared with 3Tg-iAstro ACM, returned GO terms related to extracellular matrix, focal adhesion and cadherin binding overrepresented in both WT-iAstro and 3Tg-iAstro + 4-PBA, but not in 3Tg-iAstro cells (right bottom table).

### A chemical chaperone rescues protein synthesis alterations, ER-mitochondrial interaction, and homeostatic defects of 3Tg-iAstro cells

Our data suggest that impairment of protein synthesis may represent a key feature of astrocytic dysfunction in AD, which was accompanied by the increased interaction between ER and mitochondria ^20,21^. Therefore we investigated if the correction of protein folding efficiency may be beneficial to mitigate the protein synthesis and ER-mitochondria interaction defects in 3Tg-iAstro cells using 4-PBA, an FDA approved small molecule that has shown a protective effect on the AD related neuropathology in several animal models ^35,36^. Moreover, it has been proven that 4-PBA, due to its action as a chemical chaperone, can promote correct protein trafficking, folding and prevent protein aggregation ^35^. Incubation of cell culture with 4-PBA (3 μM, 48 h), fully rescued both protein synthesis rate (Fig. 9a) and p-eIF2α levels (Fig. 9b) in 3Tg-iAstro cells. Moreover, ER-mitochondria interaction at 8-10 nm, measured with SPLICS, was significantly lower in 4-PBA-treated 3Tg-iAstro cells ^20,24^. Strikingly, when 4-PBA-treated 3Tg-iAstro cells were plated together with pericyte/EC, effects on tubulogenesis were also reinstated (Fig. 9c). Importantly, 4-PBA treatment had no effect on pericyte/EC or pericyte/EC/WT-iAstro co-cultures. These data suggest that protein synthesis alterations compromise homeostatic functions in AD astrocytes and the improvement of astrocytic protein synthesis may rescue these alterations.

**Figure 9.**
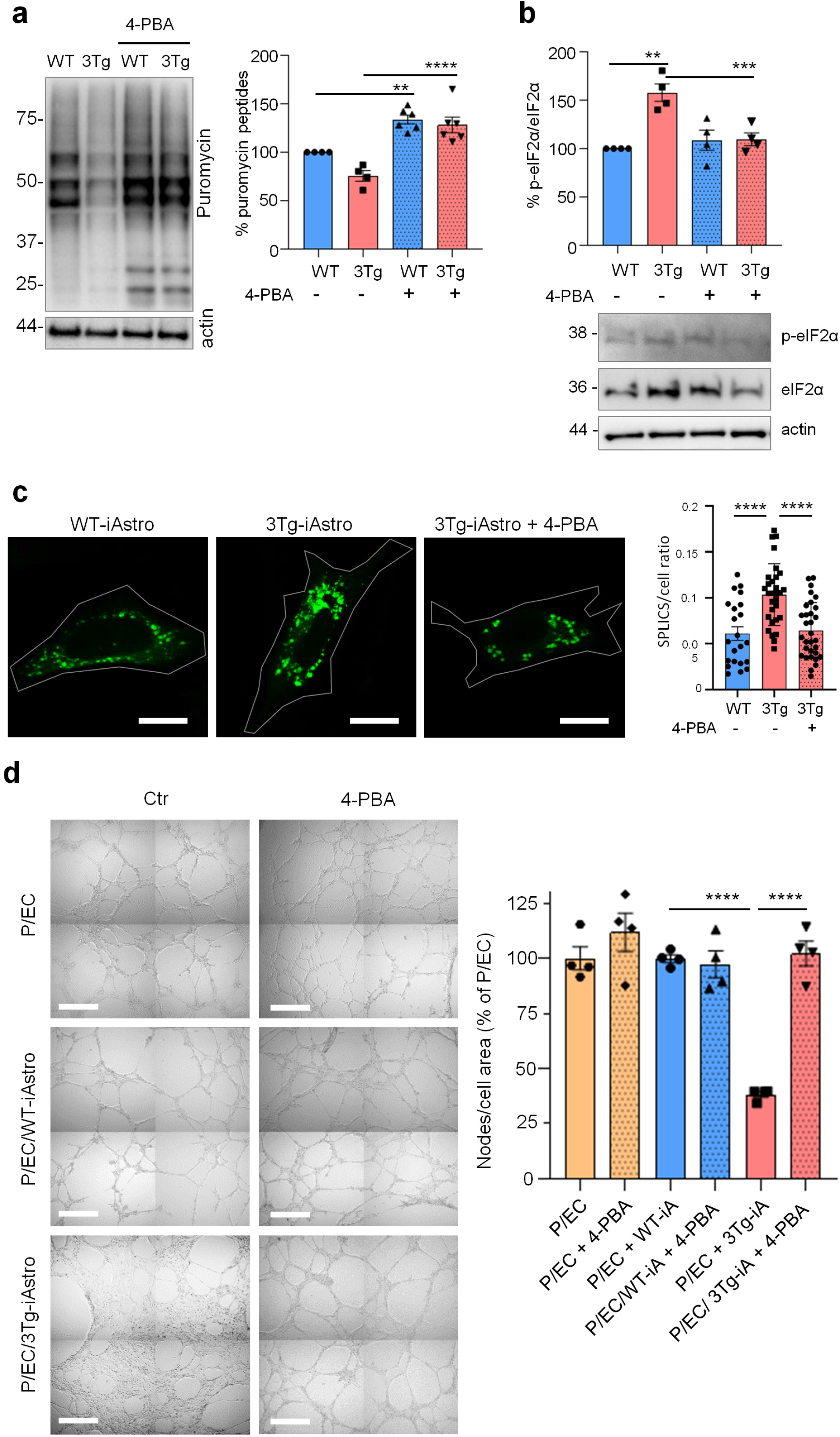
4-PBA rescues protein synthesis and p-eIF2α in 3Tg i-Astro and tubulogenesis in pericyte/EC co-cultures. **(a)** WT- and Tg-iAstro were treated or not with 4-PBA 3μM, for 48h, cells were pulsed with puromycin 4 μM and analysed by WB with anti-puromycin antibody and ponceau staining. Data are expressed as mean ± SEM from 6-4 independent experiments; **, p < 0.01; ***, p < 0.001, one-way ANOVA, Sidak’s multiple comparisons. **(b)** WB analysis of eIF2α phosphorylation on WT-iAstro and 3Tg-iAstro, treated or not with 4-PBA 3 μM, for 48h. Data are expressed as mean ± SEM from 4 independent experiments; **, p < 0.01, one-way ANOVA, Sidak’s multiple comparisons. **(c)** Representative images and quantification of SPLICS fluorescence, indicating ER-mitochondrial contacts at ~8–10 nm distance, in WT-iAstro, 3Tg-iAstro, and in 3Tg-iAstro treated with 4-PBA (3μM, for 48h). Data are expressed as mean ± SEM of n = 22 (WT-iAstro), n = 30 (Tg-iAstro), n=32 (Tg-iAstro + 4-PBA), from 3 independent coverslip, ****, p < 0.0001, one-way ANOVA, Sidak’s multiple comparisons. Scale bar = 20 μM. **(d)** Co-cultures of pericytes, endothelial cells and WT-iAstro or 3Tg-iAstro (pre-treated or not with 4-PBA for 48 h) were plated in a layer of Matrigel in presence or absence of 4-PBA (3μM). After 8 h, bright field images were taken using a Zeiss 710 confocal laser scanning microscope, scale bar = 500 μm. Data are expressed as mean ± SEM, n = 4 from 2 independent experiments; ***, p < 0.001 by one-way ANOVA, Sidak’s multiple comparisons.

**Figure 10.**
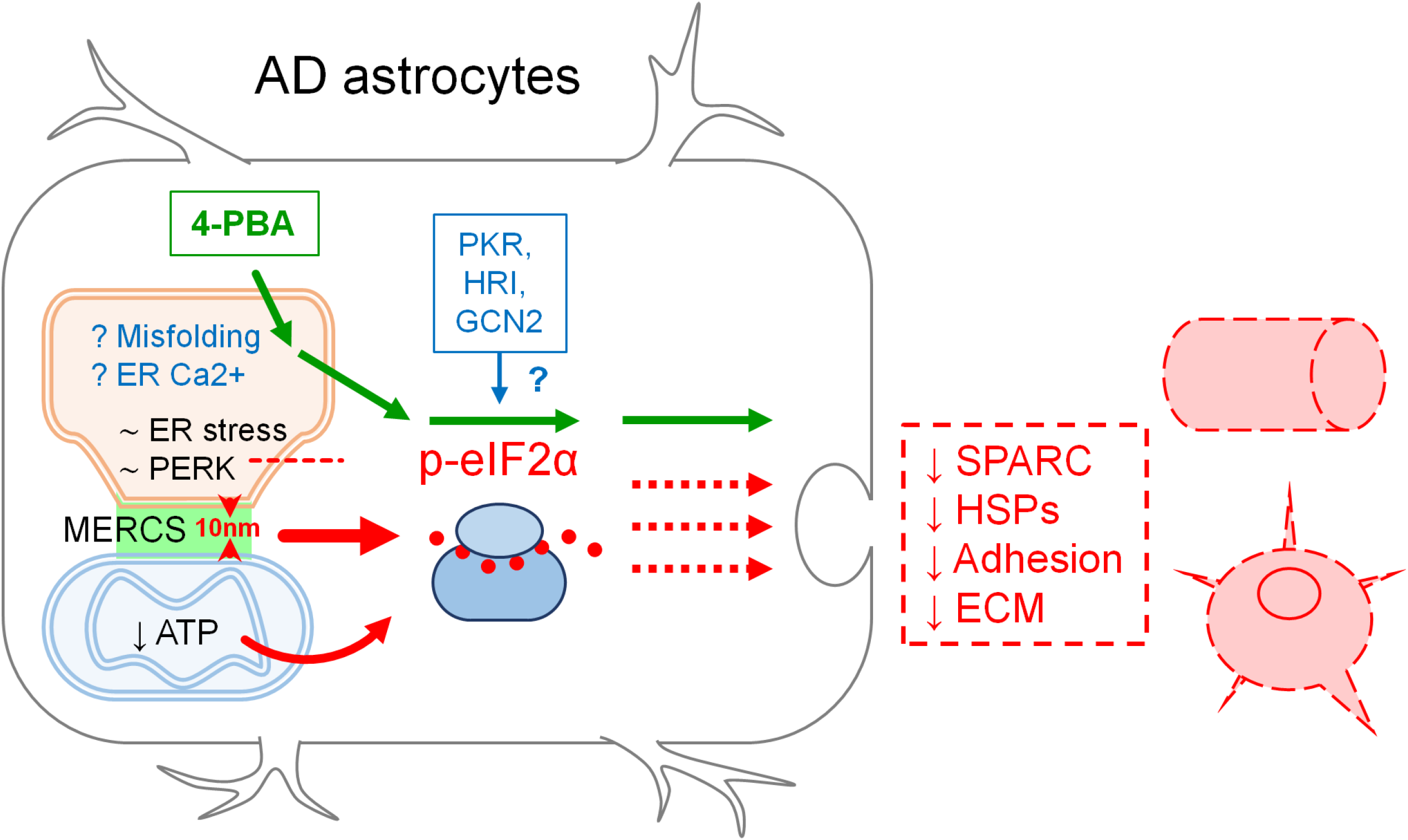
Schematic representation of the role of impaired proteins synthesis in the loss of homeostatic functions by AD astrocytes. Phosphorylation of eIF2α and reduction of protein synthesis in AD astrocytes occurs without induction of overt ER stress/UPR and activation of PERK (~ ER stress, ~ PERK and intermittent red line). A role of PKR, HRI and GCN2 kinases is to be determined (blue arrow and question mark). Alteration of ER-mitochondrial interaction could be a plausible candidate (10 nm, red arrowheads and red thick arrow), as well as a reduced ATP supply by mitochondria (curved red arrow). A role of protein misfolding and ER Ca^2+^ dyshomeostasis is hypothesized (≥ Misfolding, ≥ ER Ca^2+^). The deregulation of proteins synthesis may potentially result in impaired secretion (red intermittent arrows) of neurotrophic and neuroprotective molecules as well as impaired formation of extracellular matrix (SPARC, heat shock proteins (HSPs), Adhesion, ECM). Protein synthesis, p-eIF2α levels and homeostatic functions can be rescued by the chemical chaperone 4-PBA (green arrows).

## DISCUSSION

In the present report, we further investigated AD-related cellular alterations in a recently generated astrocytic AD model, WT- and 3Tg-iAstro cells ^19,20^. Our results suggest that in hippocampal AD astrocytes, *in vitro* and *in vivo*, the reduction of protein synthesis is driven by phosphorylation of eIF2α independently of PERK activation. Our results also suggest that p-eIF2α increase and reduction of protein synthesis may be associated with the altered ER-mitochondria interaction. We also show that 3Tg-iAstro cells exert a reduced support to primary neurons, impair pericyte/EC mediated tubulogenesis and have an altered repertoire of secreted proteins. Last, we show that improvement of protein folding in 3Tg-iAstro cells by a chemical chaperone rescued p-eIF2α levels, protein synthesis, ER-mitochondrial interaction and tubulogenesis in a three-cell pericyte/EC/astrocyte co-culture.

### PERK-independent eIF2α phosphorylation and protein synthesis inhibition in 3Tg-iAstro cells

The novel finding of this work is represented by the PERK-independent phosphorylation of eIF2α in an astrocytic cellular model of AD. Phosphorylation of eIF2α is a central switch for translation inhibition during the integrated stress response (ISR) and ER stress/UPR ^9,10^. At the present stage of knowledge four kinases are implicated in eIF2α phosphorylation during ISR, depending on the stress stimulus. PERK phosphorylates eIF2α in response to the ER accumulation of misfolded/unfolded proteins, alterations of ER Ca^2+^ levels or impairment of protein secretion machinery, which are the principal causes of ER stress ^9,37^. Protein kinase RNA-activated (PKR) has been shown to phosphorylate eIF2α in response to viral infection. General control non-derepressible 2 (GCN2) kinase is activated in response to amino acid starvation; while heme-regulated inhibitor (HRI) was initially shown to be activated by heme deprivation and to be an important component of ISR induced by oxidative or osmotic stress, heat shock, and proteasome inhibition. All these kinases have been implicated in AD pathogenesis ^7,38–40^. Therefore, further experiments are necessary to investigate their role in phosphorylation of eIF2α *in vitro* in 3Tg-iAstro cells. GADD34 is required for PP1 phosphatase to interact with and dephosphorylate PERK and is a stress-inducible protein downstream of ATF4 ^9^. Downregulation of GADD34 both in 3Tg-iAstro cells and in hippocampi of 3xTg-AD mice corroborates the conclusion on PERK-independent eIF2α phosphorylation and suggests that it may account for the increased p-eIF2α.

### Increased ER-mitochondria interaction as a cause of impaired protein synthesis in iAstro

A mounting body of evidence suggests that the ER-mitochondria interaction is increased in AD, and this may be linked to Ca^2+^ signaling deregulation, an important part of AD-related astrocytic dysfunction ^21,41–45^. Our results suggest that the impairment of protein synthesis, which is an important feature of early AD pathogenesis in both neurons and astrocytes ^46–48^, may also be caused by the augmented interaction between ER and mitochondria. The overexpression of a synthetic linker, which fixes the distance between ER and mitochondria at about 10-12 nm, faithfully reproduced the phosphorylation of eIF2α, the impairment of protein synthesis in absence of overt activation of ER stress/UPR. These results corroborate our previous finding on the increase of mitochondria-ER contact sites (MERCS) at a short distance of 8-10 nm, as measured by SPLICS sensor ^20,24^. The mechanisms whereby shortening of the ER-mitochondria distance may result in phosphorylation of eIF2α and impairment of protein synthesis remain currently unknown. It can be speculated that the disruption of a physiological distance between two organelles alters ribosomal localization and/or integrity, resulting in phosphorylation of eIF2α and impairment of assembly of the pre-initiation complex. Another mechanism may involve a reduction of ATP synthesis downstream of the impaired ER-mitochondria Ca^2+^ transfer ^20,21,49^, since protein metabolism is one of the most energy-consuming cellular activity (accounting for about 20% of overall cell energy consumption) ^50^. Further experiments are necessary to investigate how ER-mitochondria interaction may impact on the protein synthesis machinery.

### Loss of homeostatic support by 3Tg-iAstro

According to the current view on the role of astrocytic dysfunction in AD progression in terms of loss of homeostatic support, a “good” astrocytic AD cell model should provide a “bad” support to other CNS cells. This has been illustrated by us and other groups, e.g., showing that ACM from AD model mice produces dysfunction and degeneration of cultured primary neurons ^51–53^. In this frame, herein we show that, unlike WT-iAstro ACM, ACM collected from 3Tg-iAstro cells impairs protein metabolism in cultured neurons. Moreover, when plated together with pericytes and ECs in a 3D three-cell co-culture, unlike WT-iAstro, 3Tg-iAstro cells do not allow formation of tubular structures, characteristic of “angiogenic” pericyte-EC co-cultures. Astrocytes are known to secret factors supporting neurons in development and differentiation like thrombospondin 1, SPARC, Sparcl1, and lipocalin-2 ^32,54,55^. Astrocytes also secrete an array of proteins, including heat shock proteins, acting as protective factors against different stress factors, including oxidative stress ^33,56–58^. They also express components of the ECM and adhesion molecules which support cell-cell communication and cellular dynamics ^59–61^. Strikingly, neurogenic SPARC and protective HSP90 and HSP73 were significantly reduced in 3Tg-iAstro ACM compared with WT-iAstro ACM. In addition, proteins of cell adhesion and ECM were overrepresented in WT- but not in 3Tg-iAstro ACM. Therefore, the proteomic analysis of 3Tg-iAstro secretome strengthens the hypothesis of a reduced neurogenic and protective support and provides candidate molecules and signals to be further studied and tested for the development of AD therapy.

### Rescue of p-eIF2α, protein synthesis, ER-mitochondrial interaction and homeostatic support by 4-PBA

Our results suggest that a low-grade chronic ER stress, with a somewhat lower UPR response, albeit without PERK activation, might exist in 3xTg-AD astrocytes. Although this is corroborated by rescue of protein synthesis defect and p-eIF2α levels by a small chemical chaperone 4-PBA, these data are in an apparent contradiction, because activation of PERK is regarded as an obligatory step in a protein misfolding-associated UPR induction ^9^. Of note, in this regard, that a low-grade chronic ER stress is characteristic for melanoma cancer cells, and eIF2α phosphorylation may occur without ER stress ^62,63^. Furthermore, although 4-PBA is thought to act through prevention of protein aggregation in the ER, the full spectrum of its actions is not completely understood ^64^. Growing body of evidence suggest that the effect of 4-PBA can also be explained from a Ca^2+^ handling point of view. 4-PBA has been shown to rescue THG- and tunicamycin-induced ER Ca^2+^ depletion ^65^, to normalize ER-mitochondrial Ca^2+^ fluxes in the intervertebral discs nucleus pulposus cells subjected to a compression-induced ER-stress ^66^, to abolish THG-induced cytosolic Ca^2+^ signals in pancreatic acini ^67^ and to normalize cytosolic Ca^2+^ levels in 3-Chloro-1,2-propanediol (3-MCPD)-treated HEK293 cells ^68^. Moreover, 4-PBA has been shown to increase expression of SIGMA1R, a component and modulator of MERCS ^21,69^. These findings suggest that a direct or indirect action of 4-PBA on Ca^2+^ homeostasis and/or ER-mitochondrial interaction could also be hypothesized. Indeed, here we show that the increased ER-mitochondrial interaction in 3Tg-iAstro cells was fully rescued by 4-PBA. 4-PBA is an FDA approved drug and it has been shown to ameliorate cognitive performance and AD-related neuropathology in AD mouse models, holding a promise in AD therapy ^35,36^. Therefore, a more detailed investigation of the 4-PBA modulation of Ca^2+^ homeostasis and ER-mitochondria interaction is warranted.

## CONCLUSIONS

ER stress/UPR has gained much attention as a possible target for drug development in AD ^2,70–74^. However, somewhat paradoxical results and discrepancies between models and human data on the activation of components of the pathway, made the activation of ER stress/UPR in AD in its canonical form disputable ^2,8,12^. To add to the complexity of the phenomenon, ER stress/UPR in neurodegenerative diseases has mostly been studied or interpreted through the lens of a neuronal dysfunction, while for other CNS cells, in particular astrocytes, only fragmentary data are available, which makes it difficult to draw a “whole picture” ^16^. Although deregulation of protein synthesis is well documented as an early feature of AD astrocytes, the relationships between p-eIF2α, disproteostasis, and their link to ER-mitochondria communication, remain poorly understood. Our data suggest that the deregulation of protein synthesis in a model of AD astrocytes may involve p-eIF2α-associated inhibition of protein synthesis without an overt activation of PERK-mediated UPR. Herein we propose that this defective pathway may be caused by a complex array of events, including altered ER-mitochondria interaction.

## MATERIALS AND METHODS

### 3xTg-AD mice

3xTg-AD mice and non-transgenic controls (WT) were housed in the animal facility of the Università del Piemonte Orientale, with unlimited access to water and food. Animals were managed in accordance with European directive 2010/63/UE and with Italian law D.l. 26/2014. The procedures were approved by the local animal-health and ethical committee (Università del Piemonte Orientale) and were authorized by the national authority (Istituto Superiore di Sanità; authorization numbers N. 22/2013). All efforts were made to reduce the number of animals by following the 3R’s rule.

### Immortalized hippocampal astrocytes from WT and 3xTg-AD mice

Generation of immortalized astrocytes from hippocampi of WT and 3xTg-AD mice (WT- and 3Tg-iAstro cells) was described elsewhere ^19^. iAstro lines were maintained in complete culture media containing Dulbecco’s modified Eagle’s medium (DMEM; Sigma-Aldrich, Cat. D5671) supplemented with 10% fetal bovine serum (Gibco, Cat. 10270) (FBS), 2 mM L-glutamine (Sigma-Aldrich), and 1% penicillin/streptomycin solution (Sigma-Aldrich). Cells were passaged once a week and used for experiments between passages 12 and 20 from establishment.

### Pericytes and endothelial cells

Human immortalized pericytes (CL 05008-CLTH) and endothelial cells EA.hy926 (CRL-2922™) were cultured in Dulbecco’s modified Eagle’s medium (DMEM; Sigma-Aldrich, Cat. No. D5671) supplemented with 10% fetal bovine serum (Gibco, Cat. No. 10270) (FBS), 2 mM L-glutamine (Sigma-Aldrich), and 1% penicillin/streptomycin solution (Sigma-Aldrich) at 37°C in 5% CO_2_. Cells were used between passages 5 to 15 and passed twice a week.

### Hippocampal neuronal cultures

Mouse neuronal primary cultures were prepared as described previously ^17,52,75^ with slight modifications. After enzymatic and mechanical dissociation, final cellular pellet was resuspended in neurobasal A medium (Invitrogen, Cat. 10888022) supplemented with 2% B27 supplement (Invitrogen, Cat. 17504044), 2 mg/mL glutamine, 10 U/mL penicillin, and 100 mg/mL streptomycin, and plated as described above. Half of medium volume was changed every third day and the cells were lysed at days in vitro (DIV) 15.

### Cell transfection

3×10^4^ cells/well (WT- or 3Tg-iAstro) were resuspended in 250 μl of complete DMEM and 250 μl of transfection mix, and plated onto 13 mm glass coverslips in 24 well plates. For the transfection mix Lipofectamine 2000 (Thermo Fisher Scientific, Cat. 11668-019) and plasmid, in ratio 1:1, were mixed in Optimem (Gibco, Cat. 11058-021); after 3 h, transfection medium was replaced with complete medium. After 48 h, cells were washed with PBS and fixed in 4% formaldehyde (Sigma, Milan, Italy). A 10 nm ER-mitochondrial linker, which fixes the ER-mitochondrial distance at 10-12 nm, a modification of a 5 nm ER-mitochondrial linker ^27^, was a kind gift from Drs György Csordás and György Hajnóczky (Thomas Jefferson University). Generation of split-GFP contact sites sensor (SPLICS) was described elsewhere ^24,25^

### Astrocytes Conditioned Medium (ACM) preparation

For the preparation of ACM, 5×10^4^ WT-iAstro and 3Tg-iAstro cells were plated in a 6 well-plate. After 24 h the media was changed with DMEM completed with FBS, 2 mg/mL glutamine, 10 U/mL penicillin, and 100mg/mL streptomycin, or neurobasal A medium (Invitrogen, Cat. 10888022) supplemented with 2% B27 supplement (Invitrogen, Cat. 17504044), 2 mg/mL glutamine, 10 U/mL penicillin, and 100 mg/mL streptomycin. 48h later, the media were collected and centrifuged at 12,000 g, for 10 min at 4°C. ACM was stored at −80°C ^52,54^.

### Cell treatment with 4-phenylbutiric acid (4-PBA)

WT-iAstro and 3Tg-iAstro cells were plated, and after 24h were treated with 3μM 4-PBA (Sant Cruz Biotechnology, Cat. sc-232961) ^62,76^. 48h later cells were lysated and then used for WB analysis.

### Cell treatment with thapsigargin

5×10^4^ WT-iAstro or 3Tg-iAstro cells were plated in a 6MW dish. 48h later they were acutely treated with thapsigargin (Tocris, Cat. 1138) (THG). For WB analysis cells were treated with THG 1 μM, for 1 h; for RNA extraction cells were treated with THG 1 μM for 4 h ^62^.

### Pericytes/EC/astrocyte co-culture

For tubulogenesis assessment, a Matrigel synthetic extracellular matrix (Corning, Cat. 356234) was used. 96 well plates were coated with 50 μl of Matrigel, gelatinized at 37°C for 30 min. Pericytes (CL 05008-CLTH, Celther Polska, Lodz, Poland), EA.hy926 (CRL-2922, ATCC) and WT-iAstro or 3Tg-iAstro cells, in ratio 1:1:1 were resuspended in 100 μl of complete DMEM and plate on the matrix at the density of 1×10^4^ cells/well, and incubated for 8 h. Phase contrast images were acquired with a Zeiss 710 confocal laser scanning microscope.

### SUnSET for assessment of protein synthesis

Global protein synthesis rate was assessed using the Surface Sensing of Translation (SUnSET) method, as previously published ^77^. Briefly, cells were incubated with 4 μM puromycin dihydrochloride (Sigma, Cat. P8833) supplemented in normal medium at 37 °C with 5% CO_2_ for 3 h. Subsequently, cell lysates were fixed for immune fluorescence analysis or western blot analysis ^20,22^. WT and 3xTgAD mice were i.p. injectd with puromycin dihydrochloride 225 mg/Kg body weight, n = 2 WT or 3xTgAD ^78^. After 1.5 h, mice were anesthetized with i.p. injection of Zoletil (80 mg/kg) and Xylazine (45 mg/kg) and intracardially perfused with cold PBS. Brains were dissected and half of brains were used for WB analysis and the other halves of brain were post-fixed in 4% paraformaldehyde. Coronal 40 μm thick cryosections were used for immunochemical staining.

### Immunofluorescence (IF)

WT-iAstro and 3Tg-iAstro cells, grown onto 13 mm glass coverslips, were treated as previously explained. Immunofluorescence was done as follows. Cells were fixed in 4% paraformaldehyde and 4% sucrose, permeabilized (7 min in 0.1% Triton X-100 in phosphate-buffered saline (PBS)), blocked in 0,1% gelatine, and immunoprobed with an appropriate primary antibody over night at 4°C. After 3 times washing in PBS, an Alexa-conjugated secondary antibody (1:200) was applied for 1 h at room temperature (RT). The following primary antibody was used: anti-Puromycin (Millipore, Cat. MABE343). Secondary antibody was Alexa Fluor 488 anti-mouse IgG. Nuclei were counter-stained with 4’,6-diamidino-2-phenylindole (DAPI).

### Quantitative fluorescence image analysis

Images were acquired using a FV-1000 Olympus laser confocal scanning system, Zeiss 710 confocal laser scanning microscope, Leica SP8 LIGHTNING Confocal Microscope imaging systems and Leica Thunder imager 3D live cell. Images were acquired under non saturating conditions and analysed with Fiji ImageJ 1.52p software. To determine the amount of the puromycin labelled peptides on i-Astro, the puromycin mean fluorescence was measured for each selected cells excluding nucleus and expressed as fold change relative to control. To determine the amount of the puromycin labelled peptides on neuronal cultures and transfected i-Astro, the puromycin fluorescence was measured for the entire cell area excluding nucleus as a corrected total cell fluorescence (CTCF) = Integrated Density — (Area of selected cell X Mean background fluorescence). Data are expressed as fold change relative to control. For Puromycin IHC quantification, the puromycin mean fluorescence was measured by setting threshold analysis. Data are expressed as fold change relative to Ctr. For tubulogenesis assessment, the number of nodes taken with Leica Metafluor software was divided for the area covered by cells, analysed with Fiji ImageJ 1.52p software. The area covered by cells was expressed as the difference between the entire area and the closed area delimited by the tubules. Quantification of SPLICS fluorescence was performed as described elsewhere ^20^.

### Western Blot

Astroglial cultures or neuronal cultures were lysed with 100μL of lysis buffer (50mM Tris-HCl (pH 7.4), sodium dodecyl sulphate (SDS) 0.5%, 5mM EDTA, 10 μL of protease inhibitors cocktail (PIC, Millipore, Cat. 539133) and phosphatase inhibitor cocktail (Thermo Fisher Scientific, Cat. 78428) and collected in a 1.5 ml tube. Lysates were boiled at 96°C for 5 minutes and then quantified with QuantiPro BCA Assay Kit (Sigma, Cat. SLBF3463). 40 μg of proteins were mixed with the right amount of Laemmli Sample Buffer 4X (Bio-Rad), and boiled. Then samples were loaded on a 12% polyacrylamide-sodium dodecyl sulphate gel for SDS-PAGE. Proteins were transferred onto nitrocellulose membrane, using Mini Transfer Packs or Midi Transfer Packs, with Trans-Blot® Turbo ^TM^ (Bio-Rad) according to manufacturer’s instructions (Bio-Rad). The membranes were blocked in 5% skim milk (Sigma, Cat. 70166) for 45’ at room temperature. Subsequently membranes were incubated with indicated primary antibody, overnight at 4°C. Primary antibodies used are listed in Table 1, anti-β-Actin was used to normalize protein loading.

**Table1.** List of primary antibodies.

| Primary antibody | Dilution WB | Dilution ICC/IHC | Cat. N. | House |
| --- | --- | --- | --- | --- |
| Anti-ATF 4 | 1:500 |  | 390063 | Santa Cruz |
| Anti-eIF2α | 1:500 | | 133132 | Santa Cruz |
| Anti-p-eIF2α | 1:500 | 1:200 | ABP-0745 | Immunological Sciences |
| Anti-Gadd 34 | 1:550 | 1:100 | OTI2B11 | Abcam |
| Anti-GFAP |  | 1:100 | MAB-12029 | Immunological Sciences |
| Anti-PERK | 1:500 |  | C33E10 | Cell Signaling Technology |
| Anti-p-PERK | 1:500 |  | 16F8 | Cell Signaling Technology |
| Anti-Puromycin | 1:1000 | 1:200 | MABE343 | Millipore |
| Anti-β-actin | 1:2000 | | A1978 | Sigma Aldrich |

Goat anti-mouse IgG (H+L) horseradish peroxidase-conjugated secondary antibody (Bio-Rad, 1:5000; Cat. 170-6516,) and Goat anti-rabbit Igg (H+L) horseradish peroxidase-conjugated secondary antibody (Bio-Rad, 1:5000; Cat. 170-6515,) were used as secondary antibodies. Detection was carried out with SuperSignal™ West Pico/femto PLUS Chemiluminescent Substrate (Thermo Scientific), based on the chemiluminescence of luminol and developed using ChemiDoc™ Imaging System (Bio-Rad). Full length uncropped original western blots used in their manuscript are provided as a single Supplemental Material file.

### IHC

Ctr and 3xTg-AD mice were anesthetized with i.p. injection of Zoletil (80 mg/kg) and Xylazine (45 mg/kg) and intracardial perfused with cold PBS1x. Brains were dissected and sagittal sections were post-fixed in 4% paraformaldehyde in PBS1x. 40 μm slices were cut at microtome at −25°C and store at −20°C in 50% PBS1x, 25% ethylene-glycol and 25% glycerol. Free-floating staining of slices were performed. Briefly, slices were incubated with blocking solution, contains 10% serum, 1% BSA, 0,5% Triton X-100 in TBS1x, for 1 h at T room. Then, primary antibodies were applied o/n at 4°C in blocking solution. After 3 washes in TBS1x, slices were incubated with corresponding secondary antibodies for 1 h at T room, washed with TBS1x 3 times and then, counter-stained with DAPI. For co-staining, GFAP labelling was performed first and amplified with secondary antibodies plus streptavidin-Cy3 (Vector, cat. SA 1300). Then, other proteins detection was performed using corresponding primary antibodies (as p-eIF2α and GADD34) and secondary antibodies-488. Images were acquired using Leica SP8 LIGHTNING Confocal Microscope imaging systems. Images were acquired under non saturating conditions and analysed with Fiji ImageJ 1.52p software.

Primary antibodies used are anti-GFAP, Anti-p-eIF2α, Anti-Gadd 34 and Anti-Puromycin, indicated in *table 1.* Secondary antibodies are horse anti-goat biotinylated antibodies (Vector, cat. BA 9500), anti-mouse (Invitrogen, cat. A 11029) and anti-rabbit-488 antibodies (Invitrogen, cat. A32731).

### Total RNA extraction and real-time PCR

Total mRNA was extracted from 1.0×10^^6^ cells using TRIzol Lysis Reagent (Invitrogel, Cat. 15596026) according to manufacturer’s instruction. First strand of cDNA was synthesized from 0.5-1 μg of total RNA using Im-Prom-II system (Promega, Cat. A3800). Real-Time PCR was performed using iTaq qPCR master mix according to manufacturer’s instructions (Bio-Rad, Cat. 1725124) on a SFX96 Real-time system (Bio-Rad). To normalize raw real time PCR data, S18 ribosomal subunit was used. Data are expressed as delta-C (t) of gene of interest to S18 allowing appreciation of single gene expression level. Oligonucleotide primers were as follows: Atf4 (NM_009716.3), forward: GTTTAGAGCTAGGCAGTGAAG, reverse: CCTTTACACATGGAGGGATTAG; Xbp1 spliced (Xbp1s, NM_001271730.1), forward: AGTCCGCAGCAGGTG, reverse: GGTCCAACTTGTCCAGAATG; Herpud1 (NM_022331.2), forward: GTGGAGGAAGATGATGAGATAAA, reverse: CTCAGCGAGGAGTAGAAGTA; S18 (NM_011296), forward: TGCGAGTACTCAACACCAACA, reverse: CTGCTTTCCTCAACACCACA

### Proteomic analysis on astrocytes conditioned media

Astrocyte-conditioned media from WT-iAstro and 3Tg-iAstro cells, the cells treated with 4-PBA (3 μM, 48h) or expressing 10nm-EML (8-10 ml from three independently plated 10 cm Petri dish per condition) were collected, proteins were precipitated by TCA, washed and digested with trypsin. 100 μg of protein in 25 μl of 100 mM NH4HCO3 were reduced with 2.5 μL of 200 mM DTT (Sigma) at 90°C for 20 min and alkylated with 10 μL 200 mM iodoacetamide (Sigma) for 1h at RT protected from light. Any excess of iodoacetamide was removed by the addition of 200 mM DTT. The samples were then digested with 5 μg of trypsin (Promega, Sequence Grade). After an ON incubation at 37°C, 2 μL of neat formic acid were added to stop trypsin activity and the digested samples were dried by Speed Vacuum ^79^. The peptide digests were desalted on the Discovery® DSC-18 solid phase extraction (SPE) 96-well Plate (25 mg/well) (Sigma-Aldrich Inc., St. Louis, MO, USA)^80^.

LC–MS/MS analyses were performed using a micro-LC Eksigent Technologies (Dublin, USA) system with a stationary phase of a Halo Fused C18 column (0.5 × 100 mm, 2.7 μm; Eksigent Technologies, Dublin, USA). The injection volume was 4.0 μL and the oven temperature was set at 40°C. The mobile phase was a mixture of 0.1% (v/v) formic acid in water (A) and 0.1% (v/v) formic acid in acetonitrile (B), eluting at a flow-rate of 15.0 μL/min at increasing concentrations of B from 2% to 40% in 30 min. The LC system was interfaced with a 5600+ TripleTOF system (AB Sciex, Concord, Canada) equipped with a DuoSpray Ion Source. Samples were subjected to the traditional data-dependent acquisition (DDA) as previously described ^81^. The MS data were acquired with Analyst TF 1.7 (SCIEX, Concord, Canada). Three instrumental replicates for each sample were subjected to the DIA analysis ^82^. The MS files were searched using the software Mascot v. 2.4 (Matrix Science Inc., Boston, USA) using trypsin as enzyme, with 2 missed cleavages and a search tolerance of 50 ppm was specified for the peptide mass tolerance, and 0.1 Da for the MS/MS tolerance, charges of the peptides to search for were set to 2 +, 3 + and 4 +, and the search was set on monoisotopic mass and FDR at 1%. The instrument was set to ESI-QUAD-TOF and the following modifications were specified for the search: carbamidomethyl cysteines as fixed modification and oxidized methionine as variable modification. The UniProt/Swiss-Prot reviewed database containing mouse proteins (version 12/10/2018, containing 25137 sequence entries) was used.

The quantification was performed by integrating the extracted ion chromatogram of all the unique ions for a given peptide. The quantification was carried out with PeakView 2.0 and MarkerView 1.2. (Sciex, Concord, ON, Canada). Six peptides per protein and six transitions per peptide were extracted from the SWATH files. Shared peptides were excluded as well as peptides with modifications. Peptides with FDR lower than 1.0% were exported in MarkerView for the t-test.

### Statistical analysis

Statistical analysis and related graphical representations was done using GraphPad Prism v.7. A two-tailed unpaired Student’s t-test or one-way ANOVA test were used. No samples/results were excluded from the analysis. Differences were considered significant at p < 0.05.

## Author contribution

Conceptualization and data interpretation A.A.Genazzani, D.L. and L.T.; methodology, G.D., E.R., M. Manfredi, M.C., E.DG., A.G. and M.G.; software, M.G. and M. Manfredi; validation, M. Moro, B.P., E.T.,V.V.V., D.G., A. LF., E.B., S.V., G.D. and E.R.; formal analysis, M. Moro, G.D., E.R., M. Manfredi, D.L and L.T.; investigation, G.D., E.R., V.V.V., A.A.Grolla, M. Moro, D.L and L.T.; resources, A.A.Grolla, M.C., M. Manfredi, M.G. and E.DG.; data curation, V.V.V., M. Moro, G.D. and L.T; writing—original draft preparation, M. Moro, G.D., L.T and D.L.; writing—review and editing, G.D., L.T., D.L., A.A.Grolla, A.A.Genazzani; supervision, L.T., D.L. and A.A.Genazzani; project administration, L.T., A.A.Genazzani and D.L.; funding acquisition, M. Manfredi, A.A.Genazzani, and D.L. All authors have read and agreed to the published version of the manuscript.

## Acknowledgment

Advance imaging System-CAAD- Center for Translational Research on Autoimmune and Allergic Disease-Università del Piemonte Orientale “Amedeo Avogadro”, Novara, Italy. We thank Drs György Csordás and György Hajnóczky (Thomas Jefferson University) for kind donation of 10 nm ER-mitochondrial linker.

