## Supplemental Figure 1 for "Protein synthesis inhibition and loss of homeostatic functions in astrocytes from an Alzheimer’s disease mouse model: a role for ER-mitochondria interaction"

**Supplemental Figure 1 for manuscript:**

**Protein synthesis inhibition and loss of homeostatic functions in astrocytes from an Alzheimer's disease mouse model: a role for ER-mitochondria interaction.**

Laura Tapella[1*], Giulia Dematteis[1*], Marianna Moro[1], Beatrice Pistolato[1], Elisa Tonelli[1], Virginia Vita Vanella[4], Daniele Giustina[1], Aleida La Forgia[1], Elena Restelli[2$], Elettra Barberis[4], Tito Cali[5], Marisa Brini[6], Salvatore Villani[1], Erika Del Grosso[1], Mariagrazia Grilli[1], Marcello Manfredi[4], Marco Corazzari[3], Ambra A Grolla[1], Armando A Genazzani[1#], Dmitry Lim[1#]

**Affiliations**

[1] Department of Pharmaceutical Sciences, Università del Piemonte Orientale "Amedeo Avogadro", Via Bovio 6, 28100, Novara, Italy;

[2] Istituto di Ricerche Farmacologiche Mario Negri IRCCS, via Mario Negri 2, 20156, Milan, Italy;

[3] Department of Health Science (DSS), Center for Translational Research on Autoimmune and Allergic Disease (CAAD) & Interdisciplinary Research Center of Autoimmune Diseases (IRCAD), Università del Piemonte Orientale "Amedeo Avogadro";

[4] Department of Translational Medicine, Center for Translational Research on Autoimmune and Allergic Diseases (CAAD), Università del Piemonte Orientale "Amedeo Avogadro".

[5] Department of Biomedical Sciences, Neuroscience Center (PNC), Centro Studi per la Neurodegenerazione (CESNE), University of Padua, Padova, Italy.

[6] Department of Biology, University of Padua, Padova, Italy, Centro Studi per la Neurodegenerazione (CESNE)

[$] Current address: Human Technopole, Milan, Italy.

* These Authors contributed equally.

### Supplemental Figure 1

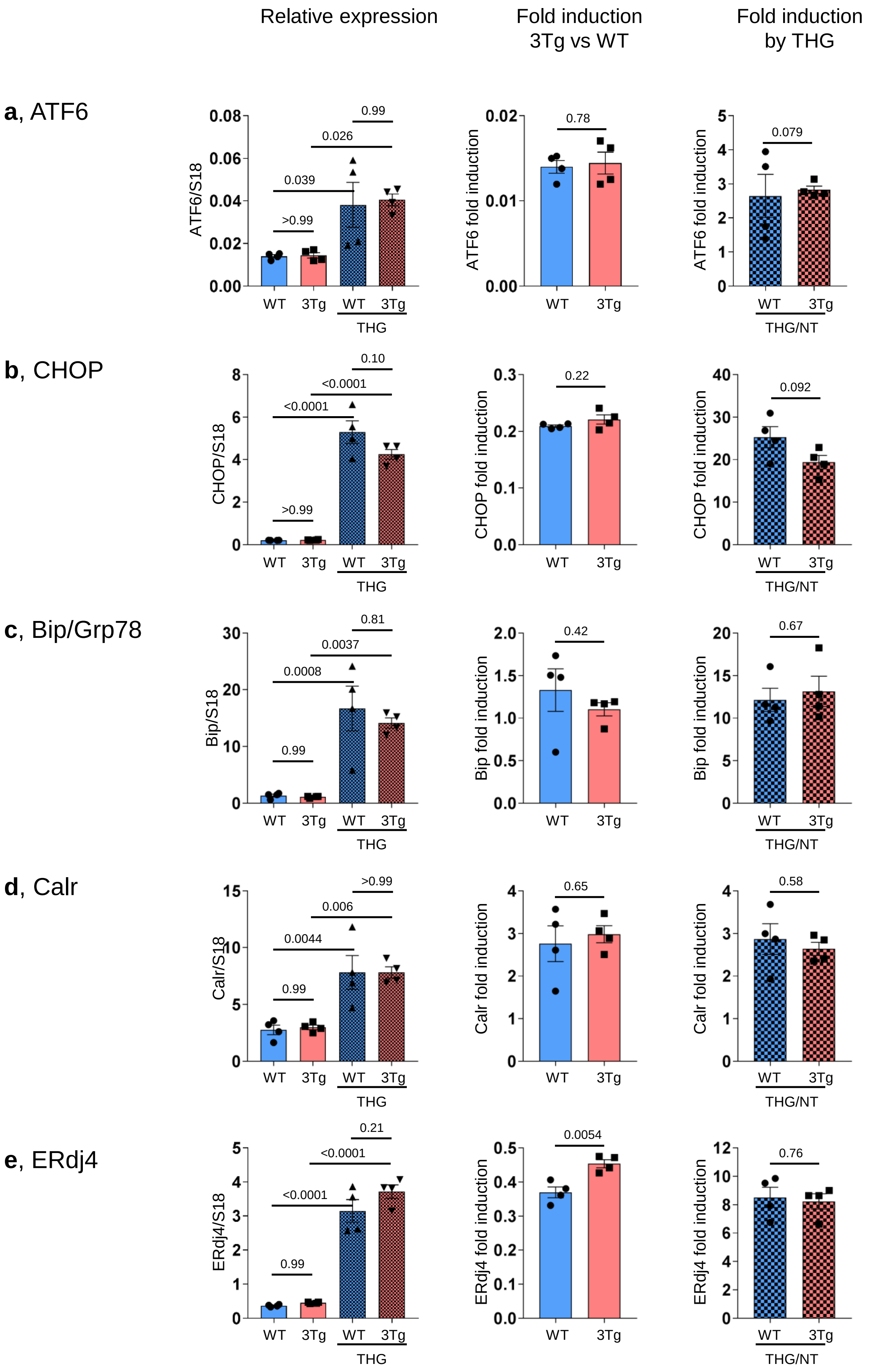

#### Supplemental Figure 1, continue.

**f**, Cartoon with ER stress/UPR pathways

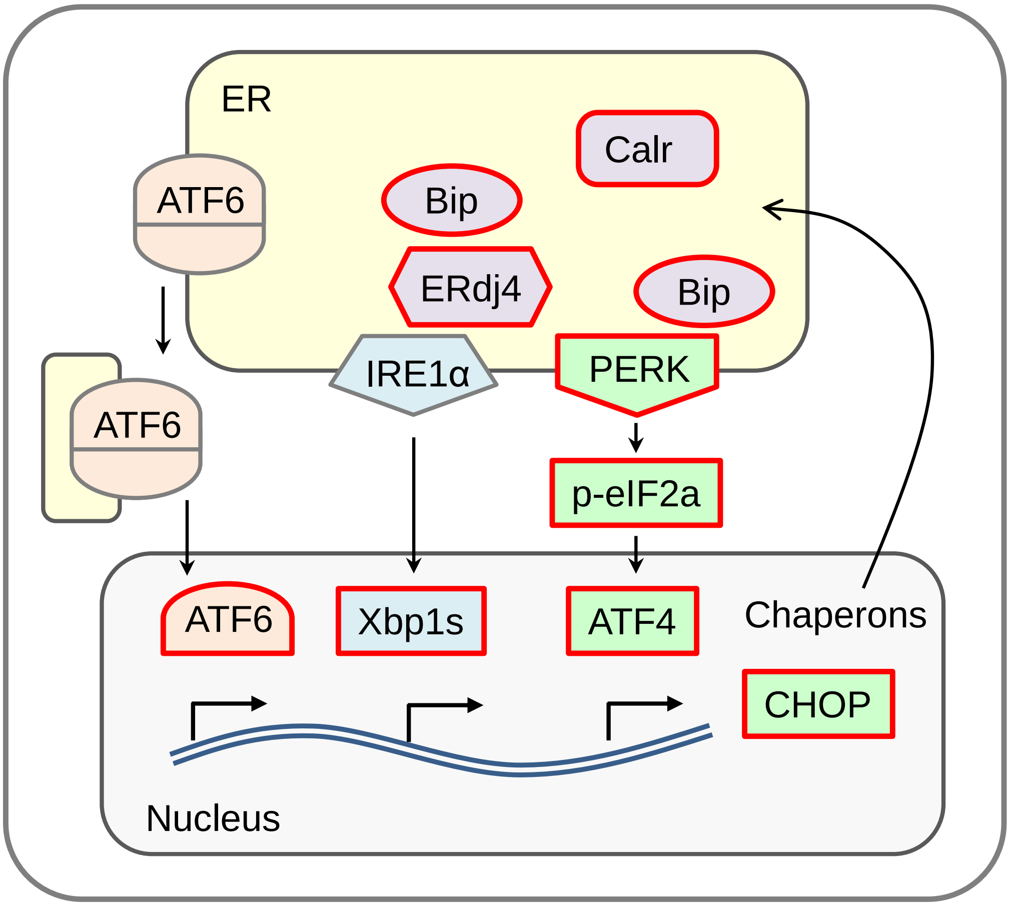


**Supplemental Figure 1. ER stress/UPR genes induction in WT- and 3Tg-iAstro cells.** Real-time PCR of Atf6 **(a)**, Chop **(b),** Bip/Grp78 (**c**), calreticulin (Calr) (**d**) and Erdj4 **(e)** transcripts in cells treated or not with 1 μM THG for 4h. Total RNA was extracted as describe in Method section, specific oligonucleotide primers were as follows (5'→3'): Atf6: forw, GATGGTGACAACCAGAAAGA, rev, TGGAGGTGGAGGCATATAA; Chop: forw, ACACGCACATCCCAAAG, rev, ACCACTCTGTTTCCGTTTC; Bip/Grp78: forw, TGCAGCAGGACATCAAGTTC, rev CTGGGGCAAATGTCTTGGTT; Calr: forw, AGGACATGCATGGAGACTCA, rev, CCGGATATCCTTGTTGATCAGC; Erdj4: forw, GGATGCCAATAGTCGGAAAG, rev, GGTGGTATCAAAGCCACTAAA. Values represent mean ± SEM ΔC(t) of gene/S18 of 4 independent experiments for each condition. Data of untreated WT- and 3Tg-iAstro cells (middle plots) and THG treated /untreated cell (right plots) are presented separately. Left plots were analyzed using ANOVA, with Tukey posthoc test; middle and right plots were analyzed using unpaired two-tail Student's t-test.
**f**, Cartoon illustrating principal routes of ER stress/UPR in which red-colored shapes indicate genes or proteins assayed in the present work.
